# Phagosomal RNA sensing through TLR8 controls susceptibility to tuberculosis

**DOI:** 10.1101/2022.06.14.496072

**Authors:** Charlotte Maserumule, Charlotte Passemar, Olivia S H Oh, Kriztina Hegyi, Karen Brown, Aaron Weimann, Adam Dinan, Sonia Davila, Catherine Klapholz, Josephine Bryant, Deepshikha Verma, Jacob Gadwa, Shivankari Krishnananthasivam, Kridakorn Vongtongsalee, Edward Kendall, Andres Trelles, Martin L Hibberd, Rafael Prados-Rosales, Kaliappan Andi, S Siva Kumar, Diane Ordway, Paul A MacAry, R. Andres Floto

## Abstract

Genetic determinants of susceptibility to *Mycobacterium tuberculosis* (*Mtb*) are poorly understood but could provide insights into critical pathways involved in infection, informing host-directed therapies and enabling risk stratification at individual and population levels. Through a genome-wide forward genetic screen, we identify the Toll-like Receptor 8 (TLR8), as a key regulator of intracellular killing of *Mtb*. Pharmacological TLR8 activation enhances killing of phylogenetically diverse clinical isolates of drug-susceptible and multidrug-resistant *Mtb* by macrophages and during *in vivo* infection in mice. TLR8 is activated by phagosomal mycobacterial RNA released by extracellular membrane vesicles, and enhances xenophagy-dependent *Mtb* killing. We find that the TLR8 variant, M1V, common in far eastern populations, enhances intracellular killing of *Mtb* through preferential signal-dependent trafficking to phagosomes. TLR8 signalling may therefore both regulate susceptibility to tuberculosis and provide novel drug targets.

**Single sentence summary:** RNA released from *Mycobacterium tuberculosis* in the macrophage phagosome is sensed by the pattern recognition receptor TLR8 controlling host susceptibility to tuberculosis and revealing a druggable pathway for host-directed therapy.

## Main text

Tuberculosis (TB), a disease caused by mycobacteria of the *Mtb* complex (MTBC), remains a major global threat to human health, with an estimated third of the World population at one time exposed (1), over 1.3 million deaths recorded per year (2), and growing rates seen for multidrug resistant (MDR) and extensively drug resistant (XDR) (3,4), Increasing antibiotic resistance and long treatment durations have motivated the search for druggable innate immune pathways that could be pharmacologically stimulated to deliver host-directed therapy (5–7).

The factors that control cell-autonomous immunity to *Mtb*, however, remain incompletely understood (1,7), despite forward genetic screens mostly using other mycobacterial species in macrophage or zebrafish infection models (8–10), genome-wide association studies in several ethnic populations (11–17) and genetic analyses of primary immunodeficiencies associated with mycobacterial susceptibility (18).

Since the initial interactions of *Mtb* with macrophages appear critical in determining the outcome of human infection (7), we set out to discover new druggable pathways in macrophages that control intracellular survival of *Mtb*.

To identify novel host restriction factors, we exposed THP1 macrophages, transduced with a genome-wide CRISPR library (19), to a GFP-expressing auxotrophic *Mtb* strain (20). We then selected cells with excess *Mtb-*associated fluorescence at 24h post infection (using Fluorescence-Activated Cell Sorting; FACS), amplified and sequenced guide RNA (gRNA) templates from extracted DNA, and then detected targeted genes that were statistically over-represented in the sorted, compared to the bulk, populations (**Figure S1**).

Our screen identified many plausible hits across a range of cellular processes, including many genes known to influence *Mtb* growth within macrophages (21–29) (**Figure 1A,B**; **Table S1-S4**). We selected novel hits within the druggable genome (30) involved in the xenobiotic response for further analysis, focusing on TLR8, a known endosomal sensor of single stranded RNA and established mediator of antiviral immunity (31). We compared the relative contribution of a panel of pattern recognition receptors (PRRs) to *Mtb* infection of primary human macrophages using siRNA knockdown (as previously (32)) and found that TLR8 silencing consistently increased viable intracellular bacteria (**Figure 1C**), an outcome phenocopied by CRISPR-mediated TLR8 knockout in THP1 macrophages (**Figure 1D**).

**Figure 1:**
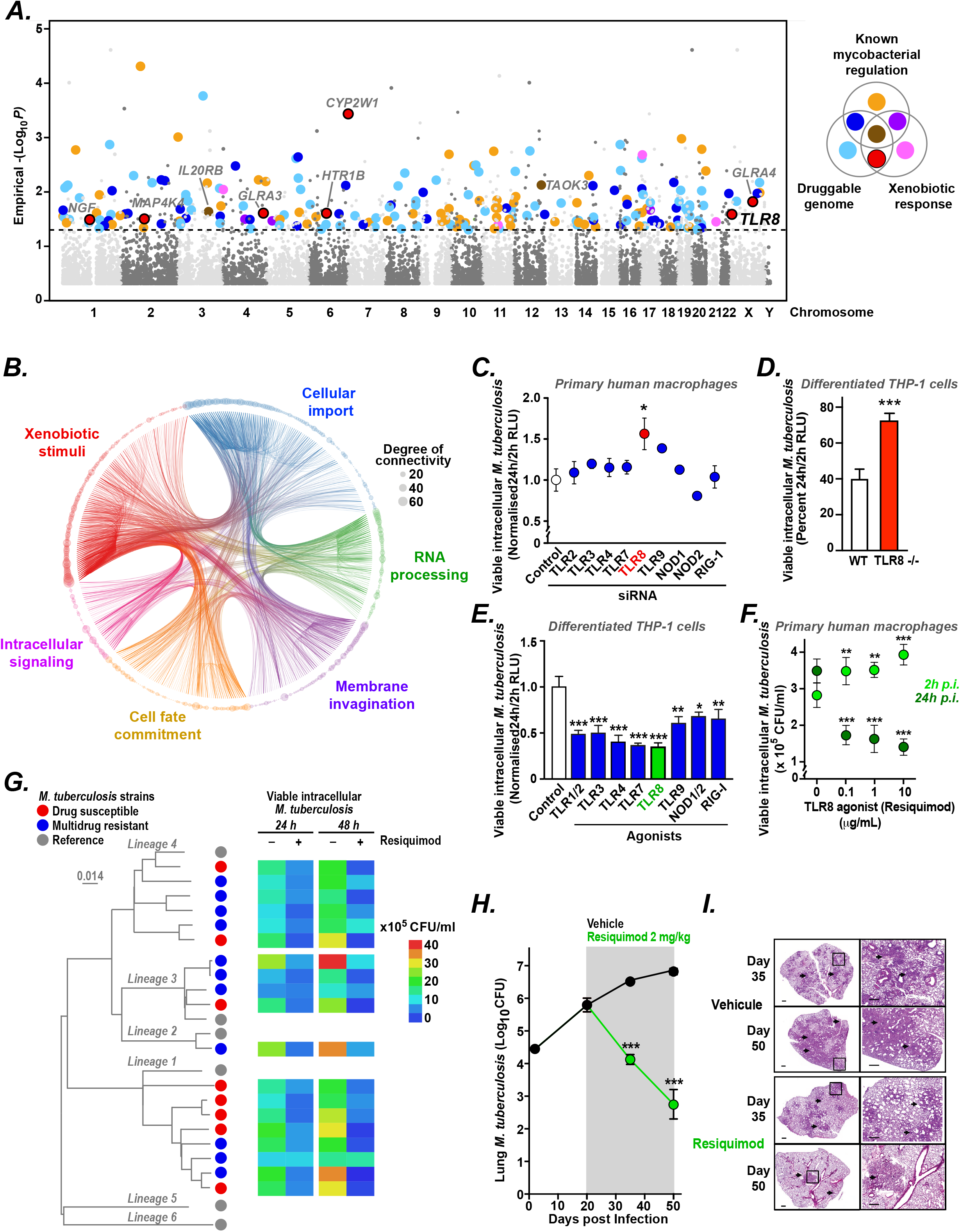
Forward genetic screening reveals TLR8 as a druggable pathway in *M. tuberculosis* infection. **(A)** Genome-wide CRISPR screen in THP1 macrophages infected with GFP-expressing *M. tuberculosis* (*Mtb*) *ΔleuD ΔpanCD* (BleuPan) performed on three separate occasions. Cells with high *Mtb*-associated fluorescence at 24 h were FACS-sorted and their gRNA representation compared to the bulk infected population at a gene level. Manhattan plot of significant hits, colour annotated based on whether genes are known regulators of mycobacterial infection, involved in the xenobiotic response, part of the druggable genome, or combinations of these. Novel druggable genes involved in the xenobiotic response (*red*) include TLR8. **(B)** Network propagation identifies interactions between gene hits in enriched GO terms (identified through REVIGO). The −log_10_(*P* value) of each gene from the CRISPR screen was used as a starting weight for propagation of the network. Node size reflects the degree of connectivity per node. **(C)** Effect on intracellular killing of luminescent *Mtb* (*H37Rv*) by primary human macrophages following knockdown (using pooled siRNA) of a panel of pattern recognition receptors (PRRs). **(D)** Intracellular killing of luminescent *Mtb* (*ΔleuD ΔpanCD* (BleuPan)) by wild type and TLR8-knockout THP1 macrophages. **(E)** Effect of agonists targeting different PRRs (*blue*), including Resiquimod targeting TLR8 (*green*), on intracellular killing of luminescent *Mtb* (*H37Rv*) by THP1 macrophages. **(F)** Effect of Resiquimod (at a range of concentrations) on intracellular killing of *Mtb* (*ΔleuD ΔpanCD* (BleuPan)) by primary human macrophages from healthy volunteers. (*C-F*) Data (mean ± SEM) shown from representative experiments at least three independent repeats, performed in at least triplicate (using primary macrophages (*C, F*) from at least three different healthy volunteers) (* p<0.05, ** p< 0.01, *** p< 0.001, Student’s t-test). **(G)** Resiquimod improves intracellular killing of clinical isolates of *Mtb*. THP1 macrophages were infected with a phylogenetically diverse collection of drug-susceptible (*red*) or multidrug-resistant (*blue*) *Mtb* clinical isolates and co-treated with Resiquimod or vehicle alone for 24 or 48 hours and viable intracellular mycobacteria were enumerated through cell-associated colony forming units (CFU/ml). Experiments were performed in at least triplicate. Maximum likelihood phylogenetic tree of all isolates tested constructed using RAxML, generated by mapping detected variable positions to *Mtb* H37Rv strain. Representatives from the main six *Mtb* lineages (grey) are included for genomic context. Scale bar indicates the number of substitutions per variable site. **(H**,**I)** Resiquimod treatment (via once daily intraperitoneal injection) of C57BL/6 mice infected with multidrug-resistant *Mtb* (TB5904) resulted in a significant reduction in lung bacterial counts (H) and improvement in lung inflammation (assessed on tissue sections treated with haematoxylin and eosin and acid-fast stains). Data represents mean ± SEM CFUs from 5 mice per time point in each group ***p<0.001 (Student t-test).

We next examined the impact of pharmacological stimulation of TLR8 and other PRRs and found that intracellular killing of *Mtb* was most enhanced by treatment with the TLR8 agonist Resiquimod (33) (**Figure 1E**), which also increased mycobacterial uptake (**Figure 1F**) but had no direct effect on *Mtb* in liquid culture (**Figure S2**). Resiquimod is an imidazoquinolinamine with antiviral and antitumour activity in preclinical animal models (34,35), clinical activity as an adjuvant to vaccines (36) and cancer immunotherapy (37), and is currently licenced by the European Medicines Agency for the treatment of cutaneous T cell lymphomas (38,39). Although Resiquimod can activate both TLR8 and TLR7 (40), its activity in *Mtb-*infected macrophages was abolished in TLR8-/-cells (**Figure S2**), supporting a specific role for TLR8 signalling. Since TLR8 in mice is a pseudogene influencing TLR7 expression (41) we were not surprised that the activity of Resiquimod on *Mtb-*infected mouse bone marrow-derived macrophages was mediated by TLR7 (**Figure S2**).

Resiquimod treatment of THP1 macrophages infected with a phylogenetically diverse collection of drug-susceptible and multidrug-resistant *Mtb* (MDR-TB) clinical isolates resulted in profound reductions in viable intracellular bacteria at 24 and 48 hours post infection (**Figure 1G**). Resiquimod treatment of mice infected via aerosol with MDR-TB isolates led to almost a three log-fold reduction in lung colony forming units (**Figure 1H, Figure S3**), with associated reduction in inflammatory lung damage (**Figure 1I**), suggesting a potential role for Resiquimod and related compounds as host-directed therapy for tuberculosis.

We next explored the mechanism of action of TLR8 during *Mtb* infection. Using a surface-expressed TLR8-TLR2 chimaeric receptor (42) stably transfected in HEK293 cells, we showed that TLR8 can be activated by a wide range of slow and rapid growing mycobacterial species (**Figure 2A**), by purified mycobacterial RNA (but not DNA) and *M. bovis* BCG lysates, and can be attenuated by RNAse pre-treatment (**Figure 2B**). During THP1 macrophage infection, TLR8 activation within *Mtb-*containing phagosomes (monitored by recruitment of MyD88) was inhibited by co-incubation with RNAse (**Figure 2C**), indicating phagosomal sensing of mycobacteria-derived RNA.

**Figure 2.**
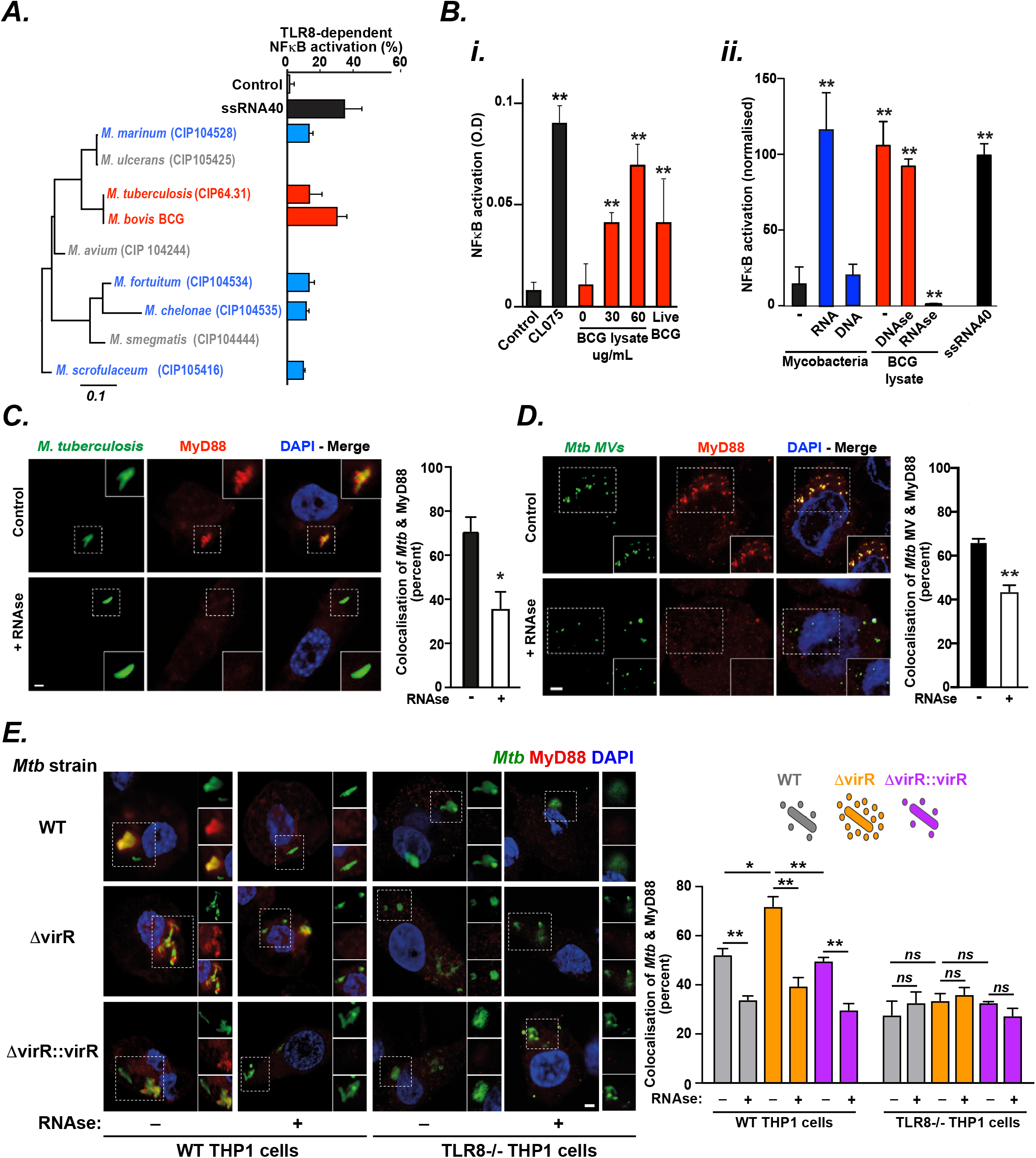
TLR8 senses phagosomal RNA released from *M. tuberculosis* in extracellular membrane vesicles. **(A)** TLR8-dependent NFκB activation in response to homogenates derived from *M. tuberculosis* and *M. bovis* BCG (*red*) as well from rapid-growing (*M. fortuitum, M. chelonae*) and slow-growing (*M. marinum, M. scrofulaceum*) nontuberculous mycobacteria (*blue*) at varying evolutionary distance from the MTB complex was assessed using NFκB reporter THP-1 macrophages (THP-1-Blue) following transfection with TLR8 siRNA or scrambled controls. The response to the TLR8 ligand ssRNA40 (*black*) is shown as a positive control. Maximum likelihood phylogenetic tree (generated using RAxML, version 2.8.2) of representative isolates shown (scale bar indicates the number of substitutions per variable site). **(B)** TLR8-2 chimaeric receptors (made by fusing the TLR8 extracellular domain to the transmembrane and cytosolic domains of TLR2) were stably surface expressed in HEK293T cells containing an NFκB luciferase reporter. TLR8-dependent NFκB signaling was assessed following (i) addition of *M. bovis* BCG lysates or live bacteria (*red*) or the TLR8 ligand CL075 (*black*) or (ii) mycobacterial RNA or DNA (*blue*), *M. bovis* BCG lystates (untreated or pre-treated with DNAase or RNAse, *red*), or ssRNA40 (*black*). (A,B) Data (mean ± SEM) are representative of at least three independent experiments performed in at least triplicate. *p< 0.05, ** p< 0.01 (Student’s t-test). **(C)** TLR8 detects *Mtb*-derived RNA within macrophage phagosomes. THP-1 macrophages were infected with GFP-labelled *Mtb* (*ΔleuD ΔpanCD* (BleuPan)) in the presence of RNAse A (100 µg/ml) or vehicle (control). Quantitative confocal microscopy was used to detect colocalization of *Mtb* (*green*) with MyD88 (*red*) in both conditions. Nuclei stained with DAPI (*blue*). **(D)** Extracellular membrane vesicles isolated from *Mtb H37Rv* were CFSE-labelled (*green; Mtb MVs*) and incubated with THP-1 macrophages either with or without RNAse A (100 μg/ml), immunostained for MyD88 (*red*), and imaged (and co-localisation quantified) using confocal microscopy. **(E)** THP-1 macrophages were infected with CFSE-labelled *Mtb H37Rv*, the transposon mutant *Tn:rv0431* (ΔVirR) which releases increased numbers of membrane vesicles, or the complemented mutant *Tn:rv0431+rv0431* (ΔVirR::VirR) in the presence or absence of RNAase A, immunostained for MyD88 (*red*), and and imaged (and co-localisation quantified) using confocal microscopy. (C-E) Image scale bar 2 μm. Data (mean ± SEM) are representative of at least three independent experiments preformed in at least triplicate (with a minimum of 50 *Mtb* phagosomes evaluated per replicate). *p< 0.05, ** p< 0.01 (Student’s t-test).

Since *Mtb* is known to produce RNA-containing extracellular membrane vesicles (MVs; **Figure S4**) (43), particularly in the context of limiting iron availability (44) (as occurs in the phagosome (45)), we wondered whether MVs might trigger TLR8 signalling. In support of this model, we found that: isolated *Mtb* MVs stimulate MyD88 recruitment to endosomal compartments within THP1 macrophages (attenuated by co-treatment with RNAse; **Figure 2D**); and that a *Mtb* mutant that over-produces MVs, ΔvirR (46,47) was able to enhance TLR8 activation (in an RNAse-inhibitable manner) during infection of wild type, but not TLR8-/-, THP1 macrophages (**Figure 2E**).

We next examined how TLR8 signaling could enhance intracellular killing of *Mtb*. Agonist stimulation of TLR8 lead to enhanced phagosome-lysosome fusion (as monitored through *Mtb* colocalization with V-ATPase; **Figure 3A**) and increased acidification (**Figure 3B**) and activity (**Figure 3C**) of lysosomes. RNA-seq analysis of the effects of TLR8 stimulation on THP1 macrophages (**Figure 3D**) revealed significant differential expression of Transcription Factor EB (TFEB) and TFEB-related genes, known regulators of lysosomal biogenesis (48) and autophagy (49). As expected, TLR8 stimulation of primary human macrophages (**Figure 3E**), and of a reconstituted heterologous expression system (**Figure S5**), led to rapid nuclear localisation of TFEB.

**Figure 3.**
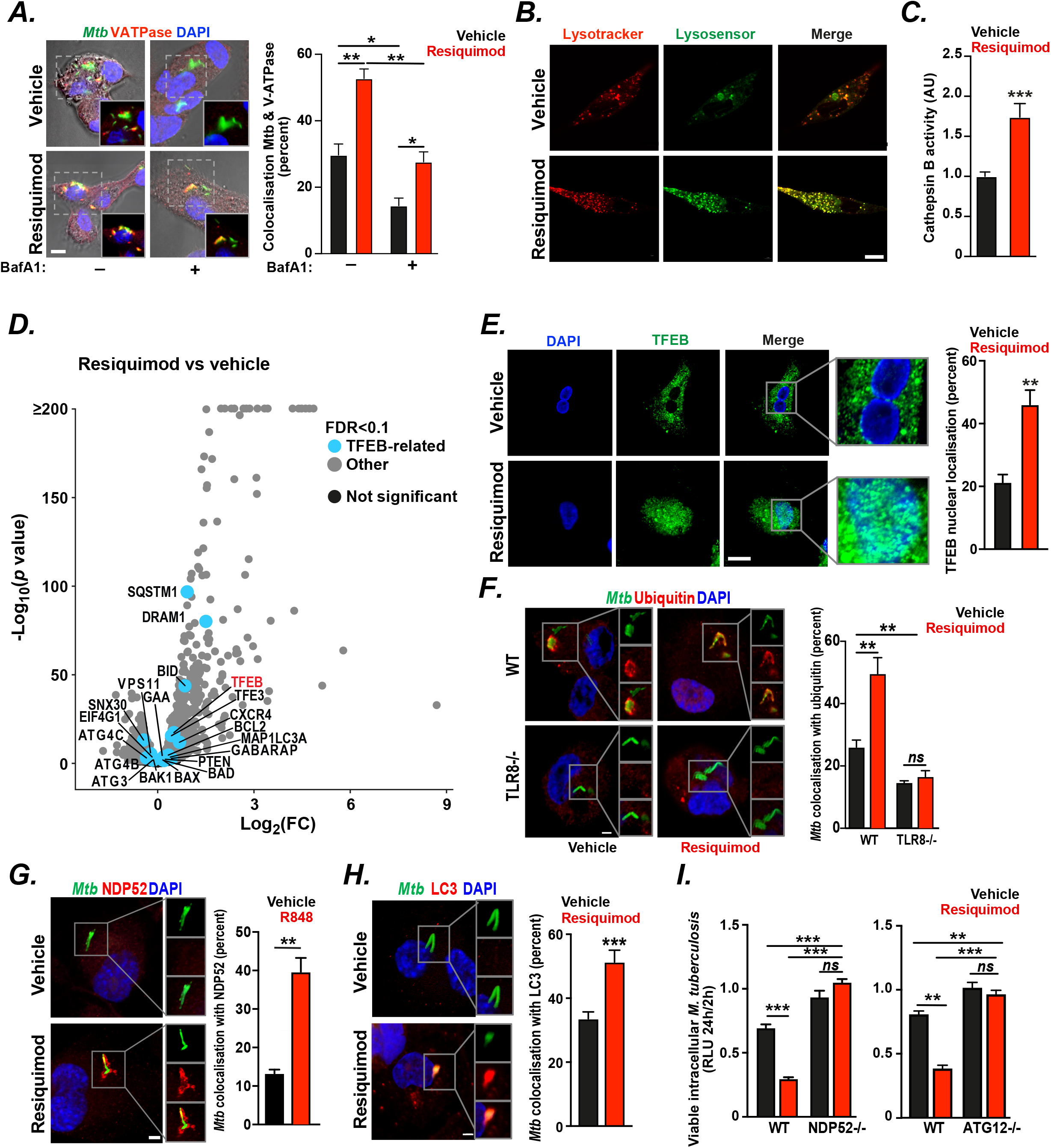
TLR8 enhances intracellular killing of mycobacteria through xenophagy. **(A)** TLR8 activation using Resiquimod increases fusion of *Mtb*-containing phagosomes with lysosomes. THP-1 macrophages infected with GFP-labelled *Mtb H37Rv* (*green*) in the presence of Bafilomycin A1 (BafA1) and/or Resiquimod (or vehicle controls) for 2 hours were immunostained for V-ATPase (*red*) and imaged (and co-localisation quantified) using confocal microscopy (scale bar 5 µm). Data (mean ± SEM) are representative of at least three independent experiments preformed in at least triplicate (with a minimum of 50 *Mtb* phagosomes evaluated per replicate). *p< 0.05, ** p< 0.01 (Student’s t-test). **(B**,**C)** Activation of TLR8 using Resiquimod increases numbers and degradative capacity of lysosomes. Uninfected THP-1 macrophages were incubated with Resiquimod and (B) subsequently incubated with LysoTrackerRed DND-99 and LysoSensorGreen DND-189 for 15 minutes prior to live confocal imaging to visualise total numbers of lysosomes (*red*) and acidified lysosomes (*green*), or (C) incubated with Magic Red™ MR-(RR)_2_ reagent for 1 hour prior to fluorescence measurements (to evaluate cathepsin B activity). Data (mean ± SEM) are representative of at least three independent experiments performed in triplicate. *** p < 0.001 (ANOVA). **(D)** Pharmacological activation of TLR8 using Resiquimod upregulates TFEB-and autophagy-related genes. Uninfected THP-1 macrophages were treated with Resiquimod for 6 hours before cell lysis and RNA extraction for RNA-seq analysis. Volcano plot shows differentially expressed genes (at a FDR (false discovery rate) < 0.1), based on three independent experiments, annotated to show TFEB (*red*), TFEB-related genes (*blue*), other significant hits (grey) and non-significant genes (*black*).**(E)** Resiquimod treatment of uninfected primary human macrophages for 24h leads to nuclear translocation (and presumed activation) of TFEB monitored and quantified by confocal microscopy (TFEB *green*, nuclei *blue*). Images and data (mean ± SEM) are representative of experiments performed in triplicate on at least three independent occasions with a minimum of 50 cells analysed per replicate. ** p < 0.01, Student’s t-test. Scale bar, 4 μm. **(F)** Wildtype (WT) or TLR8 knockout (TLR8-/-) THP-1 macrophages were infected with GFP-labelled *Mtb H37Rv* (*green*) in the presence of Resiquimod or vehicle control for 2 hours, immunostained for ubiquitin (*red*), and imaged (and co-localisation quantified) using confocal microscopy. Images and data (mean ± SEM) are representative of experiments performed in triplicate on at least three independent occasions with a minimum of 50 cells analysed per replicate. ** p < 0.01, *ns* not significant (Student’s t-test). Scale bar 2 μm. **(G**,**H)** THP-1 macrophages were infected with GFP-labelled *Mtb H37Rv* (*green*) in the presence of Resiquimod or vehicle control for 2 hours, immunostained with (G) NDP52 or (H) LC3 (*red*) and imaged (and co-localisation quantified) using confocal microscopy. Images and data (mean ± SEM) are representative of experiments performed in triplicate on at least two independent occasions with a minimum of 50 phagosomes analysed per replicate. ** p < 0.01, *** p < 0.001 (Student’s t-test) Scale bar, 2 μm. **(I)** Wildtype (WT), ATG12 knockout (ATG12-/-) or NDP52 knockout (NDP52-/-) THP-1 macrophages were infected with luminescent *Mtb* (*ΔleuD ΔpanCD* (BleuPan)) in the presence of Resiquimod or vehicle control. Viable intracellular *Mtb* at 2h and 24 h were quantified using luminescence (*RLU*). Data (mean ± SEM) are representative of experiments performed in triplicate on at least three independent occasions. ** p < 0.01, *** p < 0.001 (Student’s t-test).

We then explored the role of autophagy in TLR8 effector functions and observed: agonist-triggered increases in the number and acidification of autophagosomes in bone marrow-derived macrophages from mRFP-GFP LC3 transgenic mice (50) (**Figure S5**); agonist-induced ubiquitination of *Mtb*-containing phagosomes in wild type, but not TLR8-/-THP1 macrophages (**Figure 3F**); agonist-stimulated recruitment of the autophagy adaptor NDP52 (**Figure 3G**) and of LC3 (**Figure 3H**) to *Mtb-*containing phagosomes; and inhibition of agonist-enhanced intracellular killing of *Mtb* in NDP52-/-and ATG12-/-THP1 macrophages (**Figure 3I**). Our findings therefore indicate that TLR8 activation increases lysosomal activity and stimulates xenophagic clearance of intracellular *Mtb*.

Given the profound effects of TLR8 activation on *in vitro* and *in vivo Mtb* infections, we wondered whether naturally occurring genetic polymorphisms in *TLR8* might influence host susceptibility to *Mtb* infection through altered receptor signaling. We focused on the M1V variant (rs3764880) that leads to an alternative start codon usage and, consequently, an altered signal peptide (**Figure 4A-C**). The M1V variant has been implicated in protection from pulmonary tuberculosis in population genetic studies (41) and is found at variable allele frequencies in different ethnic groups, most abundantly in Far Eastern populations (51) (**Figure 4B**). Primary macrophages from healthy volunteers homozygous or hemizygous for the M1V TLR8 variant demonstrated enhanced killing of *Mtb* and *M. bovis* BCG (**Figure 4D**), increased inflammatory cytokine release (**Figure 4E**), and more mature mycobacteria-containing phagosomes (as evidenced by their increased size (**Figure 4F**) and greater acidification (**Figure 4G**)), compared to ancestral (wild type; WT) TLR8 controls. Transduction of M1V but not WT TLR8 into primary macrophages from WT hemizygous or homozygous individuals significantly increased phagosomal acidification (**Figure 4H**) and enhanced intracellular mycobacterial killing (**Figure 4I**), supporting a direct effect of the M1V variant on host restriction of *Mtb*.

**Figure 4:**
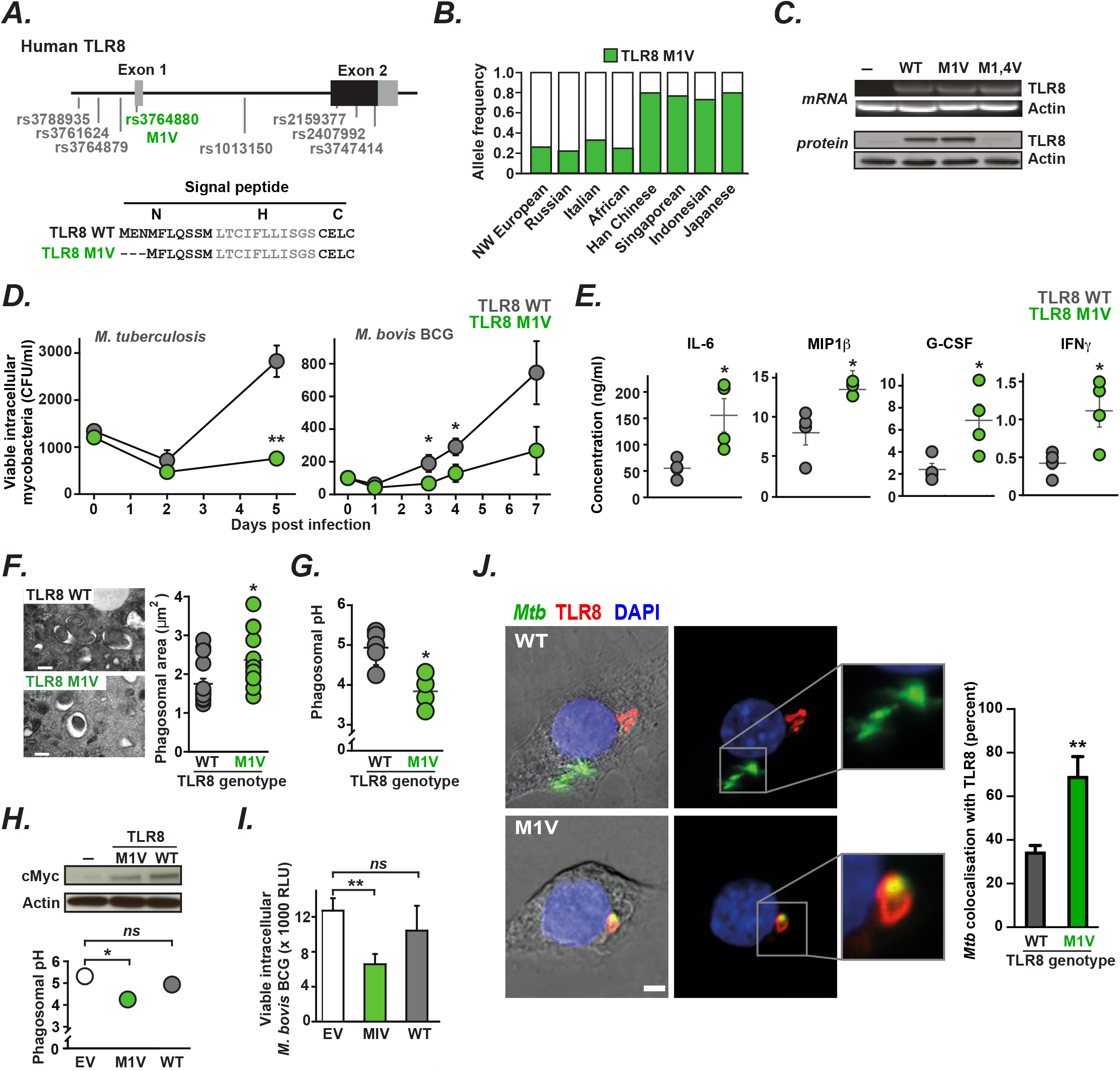
The M1V variant of TLR8 alters intracellular receptor localization and enhances intracellular killing of *M. tuberculo*sis. **(A)** (*Top*) Haplotype block of single nucleotide polymorphisms (SNPs) in human TLR8 associated with protection from pulmonary tuberculosis (43) with one non-synonymous coding polymorphism (rs3764880 M1V, *green*). (*Bottom*) Predicted signal peptide sequences of WT and M1V TLR8 suggests that the M1V polymorphism leads to alternative start codon usage (Methionine at position 4). **(B)** Allele frequency of TLR8 (*green*) in different ethnic groups (data from NCBI SNP database and (41). **(C)** RT-PCR (*top*) and western blot (*bottom*) analysis of ancestral TLR8 (WT), the M1V variant (M1V), and a mutant TLR8 with both methionines (at position 1 and 4) changed to valines (M1,4V). **(D**,**E**,**F)** Primary human macrophages from healthy volunteers that are homozygous or hemizygous for the ancestral TLR8 (TLR8 WT, *grey*) or M1V variant (TLR8 M1V, *green*) (*n* = 5 for each genotype) were infected with either *Mtb* CDC1551 or *M. bovis* BCG. (D) Viable intracellular mycobacteria were enumerated by counting CFU in cell lysates at indicated time points post infection. (E) Secreted cytokines were measure in supernatants at 24 hours post infection. (F) Mycobacteria-containing phagosomes (by electron microscopy) within primary macrophages from M1V homo/hemizygotes were larger indicating the probable formation of bactericidal phago-lysosomes. Data (mean ± SEM) are representative of experiments performed in at least triplicate. * p < 0.05, ** p < 0.01, *** p < 0.001 (Student’s t-test). (G) Primary macrophages from M1V homo/hemizygotes were able to better acidify mycobacteria-containing phagosomes than macrophages from ancestral controls, measured by assessing fluorescent ratios of internalised heat-killed *M. tuberculosis* H37Rv labelled with both acid quenchable (FITC) and pH-resistant (Alexa 633) fluorophores by flow cytometry. Data (mean ± SEM) are representative of experiments performed in at least triplicate. * p < 0.05 (Student’s t-test). **(H**,**I)** Primary human macrophages from healthy volunteers homo/hemizygous for ancestral TLR8 were transfected with either Myc-tagged TLR8 WT or M1V (or empty vector). Similar exogenous TLR8 expression was confirmed by western blot analysis using a c-Myc-specific antibody. Macrophages transfected with the M1V variant demonstrated (**H**) greater acidification of mycobacteria-containing phagosomes and (**I**) improved killing of intracellular mycobacteria. Data (mean ± SEM) are representative of experiments performed in at least triplicate on three independent occasions. * p < 0.05, ** p < 0.01 (Student’s t-test). **(J)** Mouse macrophage cells (RAW 264.7) transfected with either ancestral (WT, *top*) or M1V (M1V, *bottom*) human TLR8 tagged with c-Myc, were infected with GFP-expressing *Mtb* (*ΔleuD ΔpanCD* (BleuPan)), immunostained and imaged (and co-localisation quantified) using confocal microscopy (TLR8 *red, Mtb green*). Images and data (mean ± SEM) are representative of experiments performed in triplicate on at least three independent occasions with a minimum of 50 cells analysed per replicate. ** p < 0.01 (Student’s t-test). Scale bar 2 μm.

Since the intracellular localisation of the related receptor TLR7 is controlled by N terminal determinants (52) we wondered whether the M1V polymorphism might favourably alter TLR8 trafficking within cells. Compared to the ancestral WT receptor, we found that the M1V variant showed improved colocalization with *Mtb*-containing phagosomes when expressed in RAW-264.7 mouse macrophages (**Figure 4J**), and has an altered intracellular localisation when heterologously expressed in HEK273 cells (**Figure S6**) which is dependent on its signal peptide, since WT and M1V TLR8-CD4 chimaeric receptors are also differentially localised within cells (**Figure S6**).

In summary, we have identified TLR8 as an important mediator of cell-autonomous immunity against *Mtb* that acts by sensing mycobacterial RNA within macrophage phagosomes and stimulating xenophagic clearance. Our data suggests that TLR8 detects RNA-containing *Mtb* membrane vesicles released in response to iron starvation experienced within phagosomes (44); a mechanism that may explain the observed impact of TLR8 on macrophage responses to other bacteria (53–55).

We show that the TLR8 pathway is a potential therapeutic target during *Mtb* infection, since it is sub-maximally activated physiologically and, when stimulated pharmacologically by Resiquimod, enhances clearance of drug-susceptible and multidrug-resistant *Mtb in vitro* and *in vivo*. In addition to direct effects on macrophage clearance, TLR8 agonists may also improve adaptive immune responses during *Mtb* infection *in vivo*, since they are recognised vaccine adjuvants (36).

Finally, we demonstrate that the M1V variant that alters the signal peptide of TLR8 is preferentially trafficked to *Mtb-*containing phagosomes, and promotes greater intracellular mycobacterial killing, potentially explaining its genetic association with protection from pulmonary tuberculosis (41) and its likely evolutionary selection, and raising the possibility that polymorphisms in other genes may, singly or in combination, influence host susceptibility by regulating macrophage clearance of *Mtb*.

## Acknowledgements

We thank Dr Ben Porebski (MRC LMB, UK) for help with CRISPR screen sequencing, Dr Caetano Reis e Sousa (Crick Institute, London UK) for TLR7 mouse bone marrow, Dr David Rubinzstein for TFEB cell lines and bone marrow from mCherry-GFP LC3 transgenic mice, and Dr Brian Robertson for help with *Mtb* transfections. Primary cell in vitro experiments were authorized by regional ethics approval REC 12/ WA/0148. Funding: Supported by Wellcome Trust grants. 107032AIA (R.A.F., C.P., K.H., K.P.B., A.W., A.D.), 10224/Z/15/Z (J.M.B.), the UK Cystic Fibrosis Trust [Innovation Hub grant 001 (R.A.F., C.P., K.H., K.B., A.W., and A.D.) and Strategic Research Centre grants 002 and 010 (R.A.F D.J.O., D.V., M.J.)]; the NIHR Cambridge Biomedical Research Centre (R.A.F. and K.P.B.); Cambridge Commonswealth Trust (C.M.); Botnar Foundation grant 6063 (R.A.F., C.P., K.H., K.P.B., A.W., A.D.); NIH RO1AI162821 and Spanish MICINN contract PID2019-110240RB-I00 (R.P-R.); and the Bill and Melida Gates Foundation (P.Mc., O.S.H.O., S.K.). Author contributions: P. Mc and R.A.F. conceived the project, designed the experiments, and wrote the manuscript; C.M., C.P., O.S.H.O., K.H., K.P.B., C.K., K. A., S.S.K. performed the *in vitro* experiments; R.P.R. assisted with the membrane vesicle experiments. D.V., J.G., K.V., E.K., A.T., and D.J.O performed the mouse infection experiments; S.D. and M.L.H. performed population genetics analysis; A.W. and A.D. performed the CRISPR0-screening analyses; J.M.B performed the *Mtb* phylogenetic analysis. P.Mc. And R. A. F. provided supervisory support. Competing interests: None.

---

**Figure S1.**
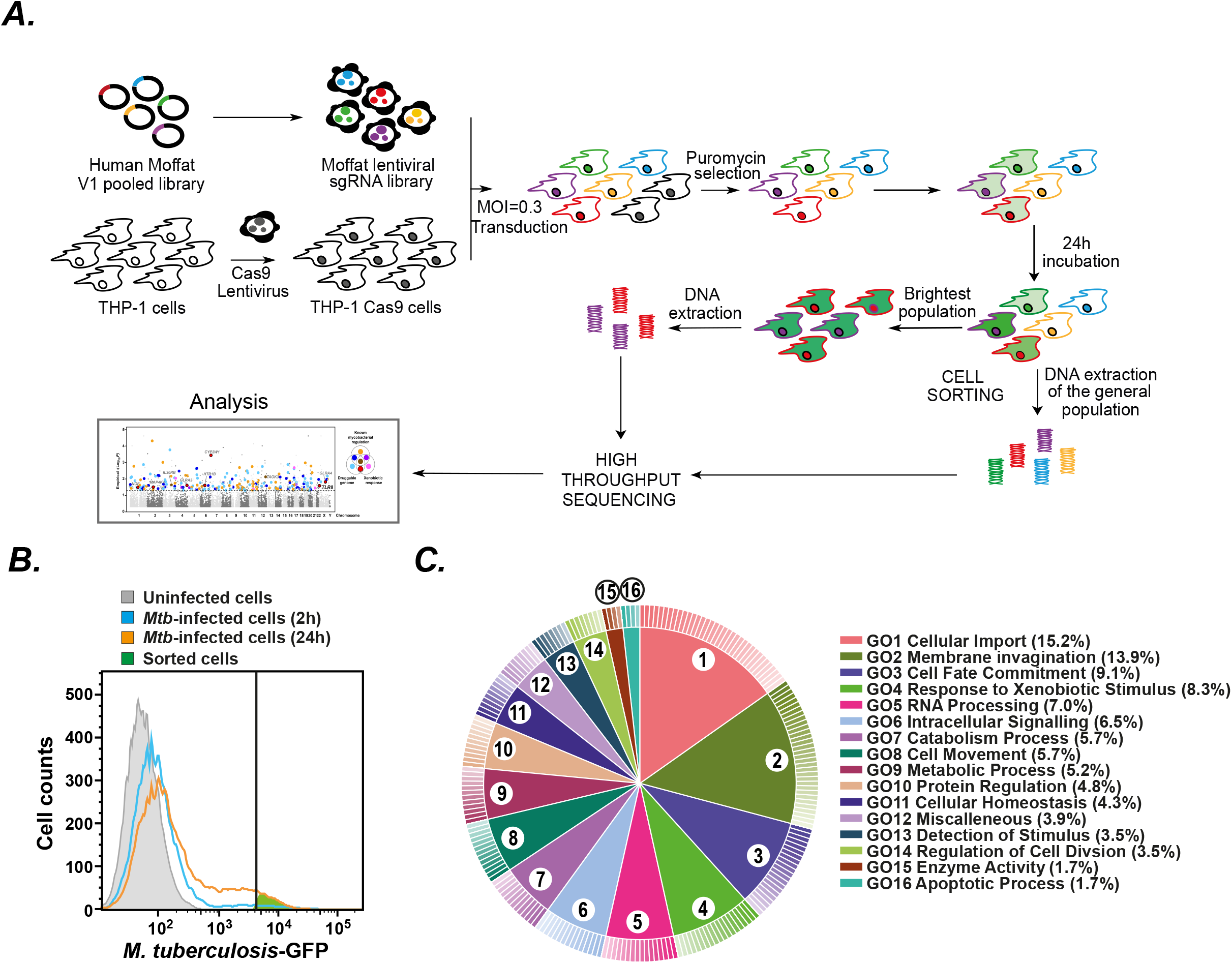
Genome-wide CRISPR screen of *M. tuberculosis-*infected THP-1 cells. (**A**) Schematic representation of CRISPR screen workflow. (**B**) Representative flow cytometry histograms of THP-1 macrophages expressing the CRISPR knockout library showing uninfected cells (grey), cell infected with GFP-expressing *M. tuberculosis* ΔleuD ΔpanCD (Bleupan) after 2 h (*blue*) and 24 h (*orange*), and the FACS-sorted population (*green*). (**C**). Visualisation of Gene Ontology (GO) terms (using CirGO (Circular Gene Ontology) python script), assigned to hits using Panther Tools and summarized and cleared for redundancy using REVIGO (Reduce & Visualize Gene Ontology terms) software.

**Figure S2.**
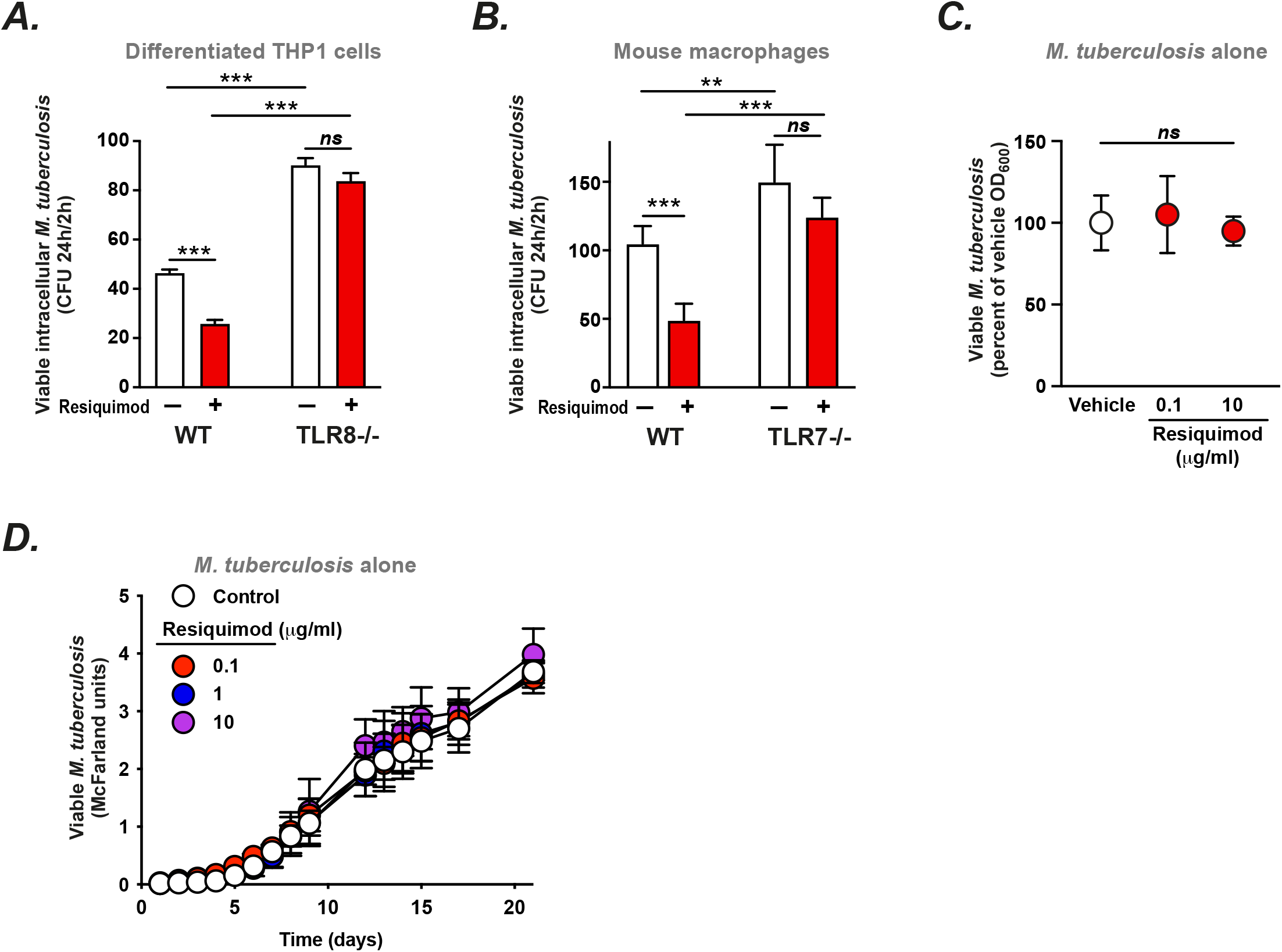
Specificity of Resiquimod for human TLR8 and mouse TLR7 in macrophages. (**A, B**) Viable intracellular *Mtb* (*Mtb* ΔleuD ΔpanCD Bleupan), expressed as the ratio of cell-associated CFUs at 24h/2h, in (**A**) wild type (WT) or TLR8 knockout (TLR8-/-) THP-1 macrophages or (**B**) bone marrow-derived macrophages from wild type (WT) or TLR7 knockout (TLR7-/-) mice, treated with Resiquimod (*red*) or vehicle alone (*white*). (**C, D**) The direct effect of Resiquimod (at a range of concentrations) on *M. tuberculosis* growth (in the absence of mammalian cells) assessed (**C**) after 24 h and (**D**) as growth curves (measured by absorbance).

**Figure S3.**
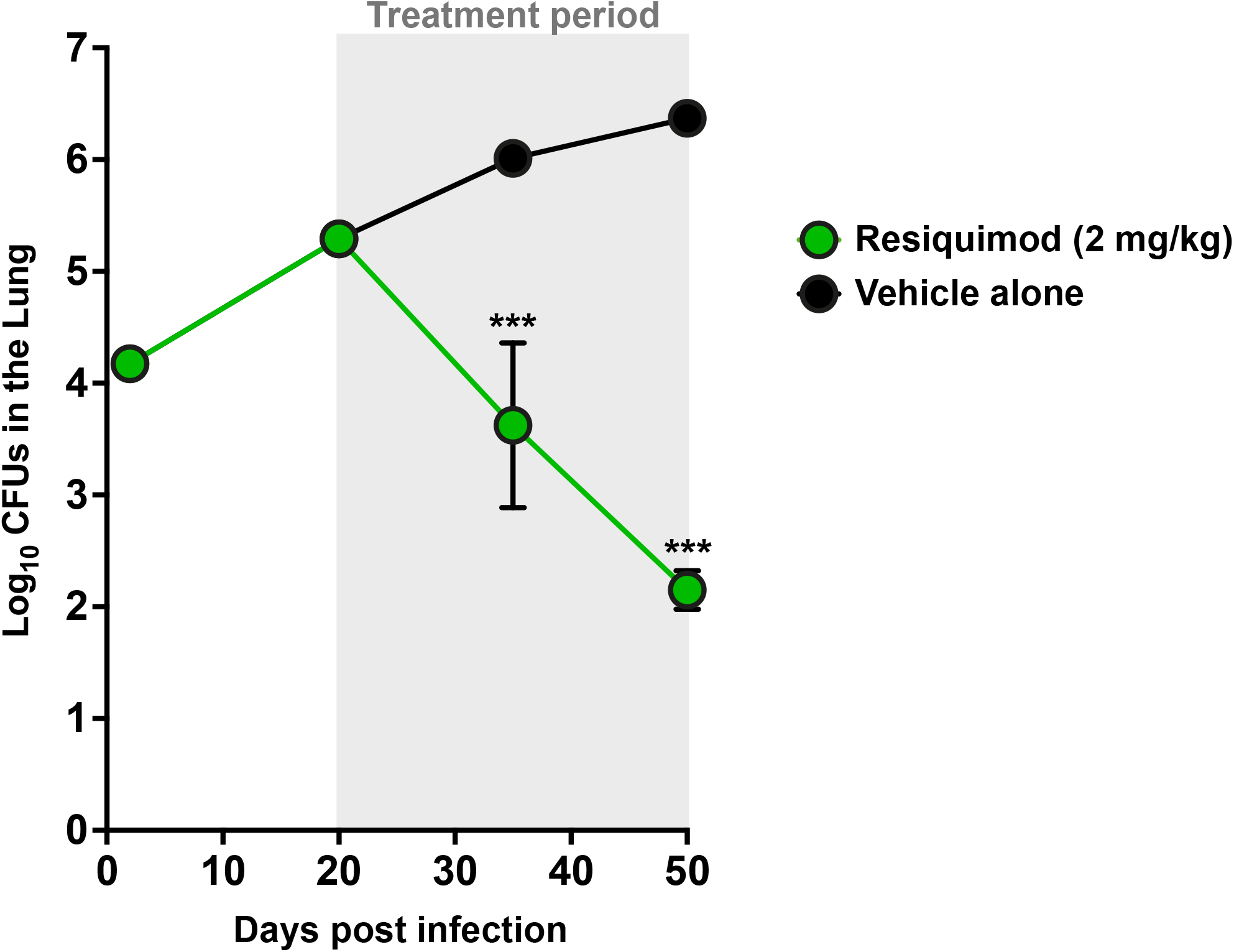
Effect of Resiquimod on *M. tuberculosis* infection *in vivo*. C57BL/6 mice were infected via aerosol with *Mycobacterium tuberculosis* MDR strain M10 and were then treated with Resiquimod (2 mg/kg, *i*.*p*. once daily; *green*) or vehicle control (*black*) for 30 days. Viable mycobacteria were enumerated at 1, 20, 35 and 50 days post-infection by plating serial dilutions of lung homogenates on nutrient 7H11 agar and quantifying CFU after 3-4 weeks incubation at 37°C, and are expressed as log10 CFU (mean ± SEM, n = 5 mice per condition). *p < 0.05, ***p< 0.001 (determined by ANOVA and the Tukey post-test).

**Figure S4.**
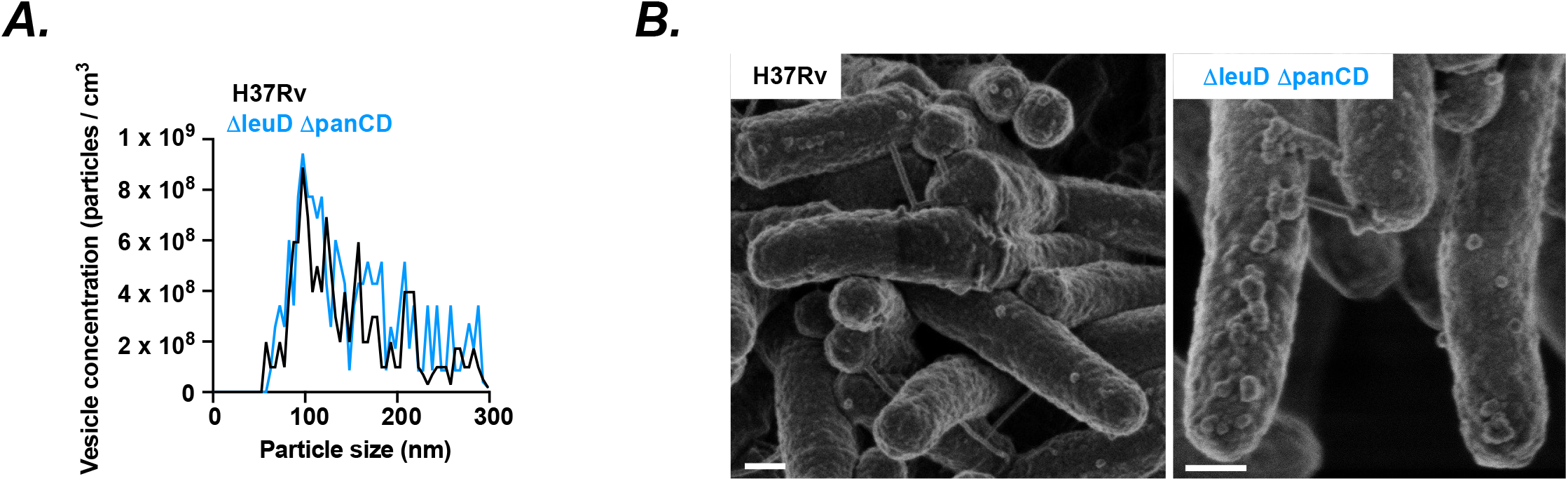
Extracellular membrane vesicles produced by H37Rv and ΔleuD ΔpanCD (Bleupan) *M. tuberculosis*. **(A)** Quantification of extracellular membrane vesicles (MV) in culture supernatants of H37Rv (*black*) and ΔleuD ΔpanCD (Bleupan) (*blue*) *M. tuberculosis* (*Mtb*) strains using nanoparticle tracking analysis. Data shown indicates mean values of three independent biological replicates, show no significant difference in MV characteristics between *Mtb* strains. (**B**) Scanning electron micrographs of H37Rv (*left*) and ΔleuD ΔpanCD (Bleupan) (*right*) *Mtb* strains illustrating MV production. Scale bar 200 nm

**Figure S5.**
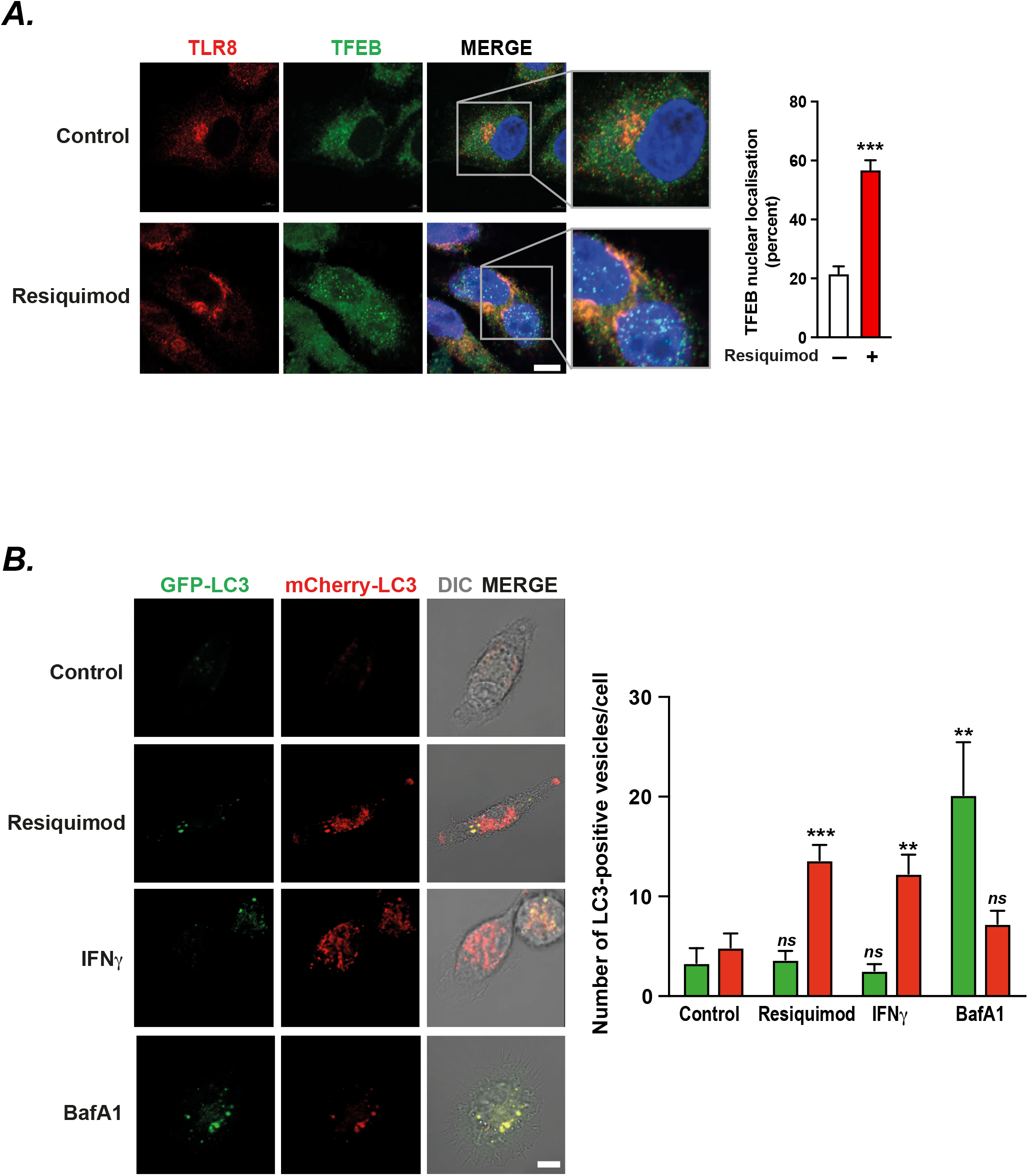
Activation of TLR8 increases TFEB translocation and stimulates xenophagy. (**A**). HeLa cells stably expressing Flag-tagged TFEB (*green*) were transfected with myc-tagged TLR8 (*red*) and treated with Resiquimod or vehicle control for 24 h. Nuclear translocation of TFEB was quantified by confocal microscopy of DAPI-stained cells (*blue*). Experiments were performed in triplicate, with at least 50 cells analysed per sample. Data (mean ± SEM) are representative of at least two independent experiments. ***p < 0.001 (Student’s t-test). Scale bar 5 μm. (**B**) Bone marrow-derived macrophages from transgenic mice expressing mRFP-GFP-tagged LC3 (56) were treated with vehicle control, Resiquimod (10 μg/ml), and interferon gamma (IFN-γ; 200 ng/ml) as a positive control for autophagic flux enhancement, or Bafilomycin A1 (BafA1; 200 nM) as an inhibitor of autophagic flux, for 24 h and imaged using live cell confocal microscopy to quantify acidified (mCherry^+^ GFP^−^; *red*) and non-acidified (mCherry^+^ GFP^+^, *green*) vesicles. Experiments were performed in duplicate, with >50 cells analysed per sample. Data (mean SEM) are representative of at least three independent repeats. **p < 0.01; ns, non-significant (Student’s t-test). Scale bar 5μm.

**Figure S6.**
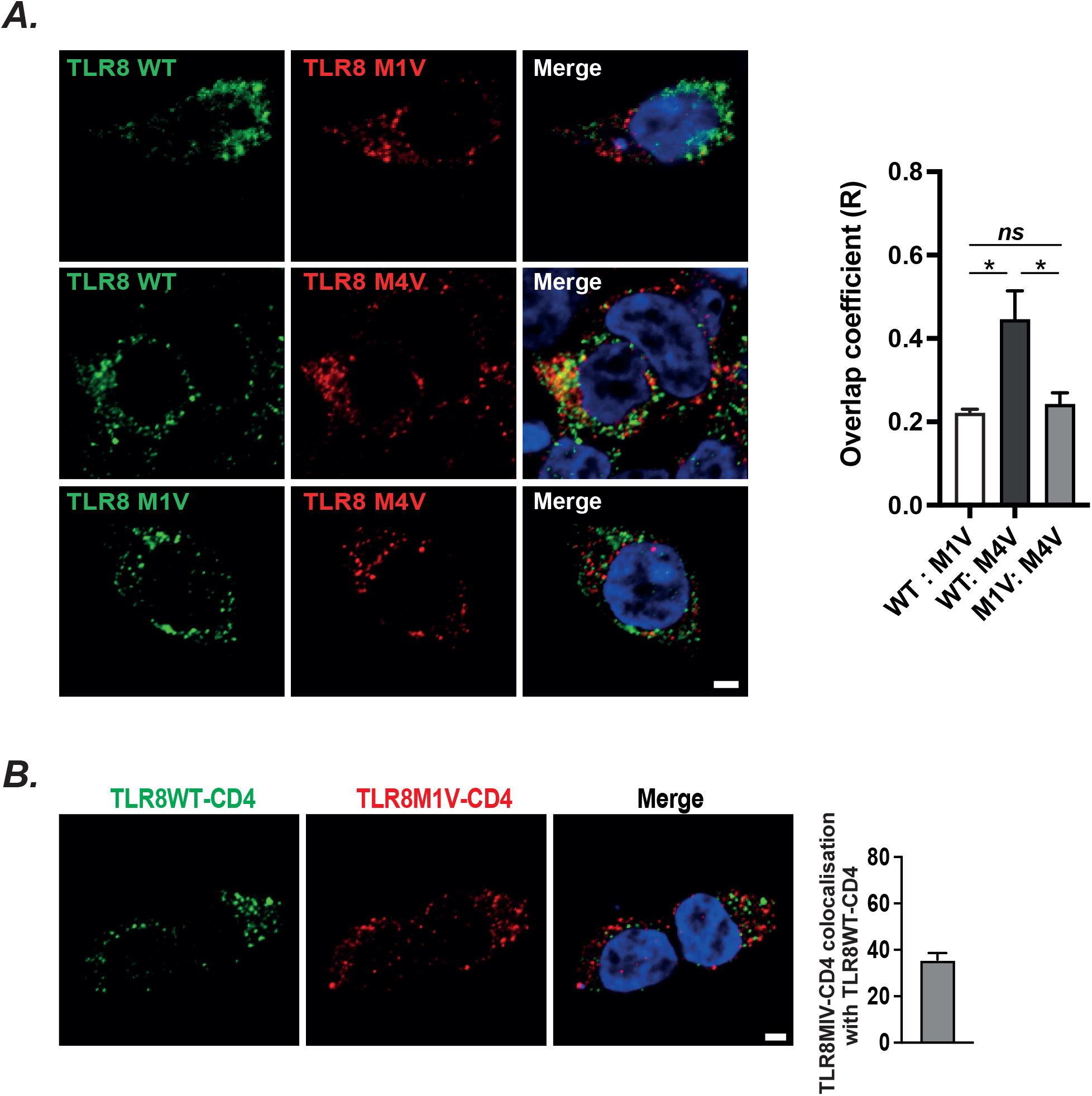
Distinct intracellular localization of ancestral and M1V variants of TLR8. (**A**) Confocal microscopy of HEK 293T cells co-transfected with two of the following TLR8 receptors: (*top*) wild type (*green*) and M1V *(red*); (*middle*) wild type (*green*) and M4V (where the second methionine has been mutated to a valine to enforce transcription from the first methionine, *red*); and (*bottom*) M1V (*green*) and M4V (*red*). Quantification of receptor co-localisation indicates only limited colocalization of the M1V variant with the other receptors, suggesting a descrete effect of the truncated signal peptide on intracellular localization. Scale bar 2 μm. (**B**) HEK293T cells were co-transfected with chimaeric receptors made from the signal peptide and intracellular domain of TLR8 joined to the extracellular and transmembrane domains of CD4. Cells were simultaneously transfected with tagged chimaeric receptors containing the wild type signal peptide (WT TLR8-CD4, *green*) and M1V signal peptide (M1V TLR8-CD4, *red*), and imaged using confocal microscopy. Quantification showing limited co-localisation of WT and M1V fusion receptors shown. Scale bar 2μm. All experiments were performed in triplicate, with at least 50 cells analysed per coverslip. Data (mean ± SEM) are representative of at least three independent experiments. Vesicles per cell and overlap coefficients were quantified using Image J. *p < 0.05, (Student’s t-test).

## Supplementary Materials

Materials and Methods

Table S1-S5

Figures S1-S5

Figure Legends

## Supplementary Materials for

**Other Supplementary Materials for this manuscript include the following**

Supplementary Tables 1-5 (Excel)

## Materials and Methods

### Antibodies

#### The following antibodies were used

MyD88 (ab2064; *Abcam*), TLR8 (HPA001608; *Sigma-Aldrich*), V-ATPase (Sc-28801; *Santa Cruz*), ATP6V1A antibody (ab137574; *Abcam*), LC3 (02316F10; *Nanotools*), NDP52 (ab124372; *Abcam*), TFEB (H00007942_M01; *Abnova*), FLAG (F1804; *Sigma-Aldrich*), Ubiquitin (ST1200; *Sigma-Aldrich*). c-myc (ab32072; *Abcam*), V5-tag (NB600-381; *Novus Biologics*), and secondary fluorescently-tagged antibodies (A11008/A11029, A21428/A32727, A21236/A21245; *Thermo Fisher Scientific*).

### Mycobacteria

#### The following strains of mycobacteria were used

*Mycobacterium tuberculosis H37Rv* (ATCC), *M. tuberculosis H37Rv-GFP* (gift from Dr Sandra Newton, Imperial London, UK*); M. tuberculosis VirR-* (*Tn:rv0431*) and its complemented strain *M. tuberculosis VirR-::WT* (*Tn:rv0431::rv0431*) (1), *M. bovis BCG*, BCG-lux, a luminescent reporter strain of *M. bovis BCG* encoding the Vibrio *lux AB* gen*e (M*.*bovis BCG-pSMT1-LuxAB)* or GFP (*M. BCG-pSMT1-LuxAB-GFP*) *(2)*, clinical isolates of *M. scrofulaceum, M. marinum, M. chelonae, M. fortuitum* (Department of Microbiology, The Yong Loo Lin School of Medicine, National University of Singapore, Singapore), multi-drug resistant (MDR) isolates of *M. tuberculosis* (TN5904 and M10) provided by Dr B. N. Kreiswirth (Public Health Research Institute TB Center, Newark, NJ) and Dr. Chan, (*Seoul, Korea*); the following clinical isolates of *M. tuberculosis* that were drug-susceptible (NS001, NS007, NS006, NS008, NS009, NS0013, NS016, NS017, NS046, NS043, NS058, NS092, NC096, NS055, NS057) and multidrug-resistant (NR006, NR007, NR009, NR004, NR010, NR008, NR021, NR024, NR029, NR033, NR041, NR045) from the Bacteriology Division of the ICMR-National Institute for Research in Tuberculosis (Chennai, India). Isolates were grown as previously described (3,4) in Middlebrook 7H9 broth (*Difco*) containing 0.5% glycerol (*Sigma-Aldrich*), 0.05% Tween 80 (*Fischer Scientific*) and 10% albumin–dextrose–catalase (ADC, *Sigma-Aldrich*) enrichment.

#### Auxotrophic M. tuberculosis

*M. tuberculosis ΔleuD ΔpanCD* (Bleupan) double auxotroph strain (5) (gift from Dr Bill Jacobs) was transduced with *pSMT12-mCherry* or *pSMT12-GFP* (gifts from Dr Lalita Ramakrishnan, Cambridge, UK) or *pSMT1-LuxAB-GFP*, and grown in Middlebrook 7H9 broth containing 0.5% glycerol, 0.05% Tween 80 and 10% oleic acid– albumin–dextrose–catalase enrichment (OADC) (BD Bioscience), 0.05 mg/ml L-leucine (*Sigma-Aldrich*), 0.024 mg/ml calcium pantothenate (Sigma-Aldrich). When necessary, 50 μg/ml hygromycin B (*EMD Millipore*), 40 μg/ml kanamycin or 50 μg/mL zeocin (*Alfa Aesar*) were added to cultures. Bacteria were grown for 15 days at 37°C, then transferred in bigger culture volume (1/100 dilution) for 10 more days in media of the same composition.

#### Mycobacterial homogenates

For generation of mycobacteria homogenates, mycobacterial cultures were harvested, washed, and resuspended in phosphate-buffered saline (PBS). Bacteria were disrupted by bead-beating in a bullet blender (Next Advance) for 5 min and homogenates were briefly centrifuged to remove the beads and intact cells. Experiments to identify the mycobacterial ligand for TLR8 were performed on *M. bovis* BCG homogenates either heat denatured at 95°C for 5min; or subjected to enzymatic digestion by RNase A (*Qiagen*) or DNase I (*Sigma-Aldrich*) for 15 minutew at room temperature. 1 unit of RNase or DNase was used for every 1μg of *M. bovis* BCG homogenate. RNA and DNA from *M bovis* BCG were obtained from cultures grown to mid-log phase using the Roche High Pure RNA Isolation kit and the DNeasy Blood and Tissue Kit (*Qiagen*) respectively, according to manufacturers’ instructions.

#### Extracellular membrane vesicles

Extracellular membrane vesicles (MVs) were isolated from *M. tuberculosis* as previously described (6). Briefly, *M. tuberculosis* H37Rv bacterial cultures were grown in 7H9 medium for 7 days and subsequently inoculated into minimal medium and incubated at 37°C for 14 days. Bacterial cultures were sequentially filtered through 0.45-μm and 0.22-μm-pore size filters, and concentrated using an Amicon Ultrafiltration system with a 100-kDa-exclusion filter. The recovered concentrate was centrifuged to recover the vesicle pellet. The membrane vesicles were purified by density gradient ultracentrifugation using OptiPrep® solution (*Sigma–Aldrich*) prepared in Dulbecco’s phosphate buffered saline (*Sigma–Aldrich*). To assess MyD88 signaling, CFSE-labelled MVs suspension in DMEM with 10% FCS were added to differentiated THP-1 cells in the presence or absence of 100 μg/ml RNAse A (*Qiagen*) and incubated at 37°C for 2 hours. The cells were then washed, fixed and immunostained for MyD88.

#### Single cell bacterial suspensions

To prepare single cell suspensions of bacteria prior to infection, bacteria were centrifuged 24 hours prior to experiment and resuspended in bacterial growth media without tween to allow one generation time and complete reformation of mycobacterial cell wall. On the day of infection, mycobacterial cultures were passed through a 27-gauge needle 10 to 12 times prior to injecting through a 5-micron filter to achieve close to single cell suspensions of bacteria.

#### Colony forming units

To enumerate colony forming units (CFU) counts, bacterial suspensions were plated on Middlebrook 7H11 agar (*Sigma-Aldrich*) with 10% OADC enrichment supplement with indicated supplements and antibiotics and CFU were counted after 21 days of incubation at 37 °C.

### Mammalian Cell cultures

#### Monocyte-derived macrophages

Peripheral blood mononuclear cells (PBMCs) were generated as previously described (3). Briefly, PBMCs were isolated from peripheral blood obtained from healthy consented subjects (approved by Regional NHS Research Ethics Committee) by Ficoll-Hypaque (*Amersham*) density separation. CD14^+^ positive selection using magnetic beads (*Miltenyi*) was used to extract monocytes, which were subsequently differentiated into macrophages by stimulation with 200 ng/ml M-CSF (*Peprotech*) in DMEM (*Sigma-Aldrich*) containing 2 mM L-glutamine (*Sigma-Aldrich*), 10% FCS (*<u>Pan Biotech</u>*), 100 U/ml penicillin/streptomycin (*Sigma-Aldrich*). Cells were differentiated for six days before assaying.

#### Bone-marrow-derived macrophages

Generation and culture of bone-marrow-derived macrophages (BMDM) was caried out as previous described (7). Femurs from 15-week-old female TLR7 knockout mice (generated as previously described (8)) or age and sex-matched C57BL/6 mouse controls were dissected to remove both ends and flushed with a 21-guage needle into serum-free DMEM. Cells were then centrifuged to remove cell culture medium and cultured in 10ml DMEM supplemented with 20% FCS, 100U/mL penicillin/streptomycin, 50μM tissue-culture grade β-Mercaptoethanol (*Sigma-Aldrich*) and 200 nM murine MCS-F (*Peptrotech*) for 3 days, after which fresh medium was added. Cells were then incubated for another 3 days before the cells were scraped and seeded onto 24-well plates for experiments.

#### THP-1 macrophages

THP-1 cells (ATCC), THP-1 BLUE NF-κB and AP-1 reporter monocytes (*Invivogen*) were maintained in RPMI 1640 (*Sigma-Aldrich*), 10% FCS, 100 U/ml penicillin/streptomycin, 2 mM L-glutamine, and 200 μg/ml Zeocin when needed. Cells were supplemented with 40 ng/mL 12-phorbol 13-myristate acetate PMA (*Peprotech*) for 48h to stimulate differentiation into macrophages.

#### Other cell lines

HeLa cells (ATCC), FLAG-tagged TFEB reporter Hela cells (gift from Dr David Rubinzstein, Cambridge, UK (9)), HEK 293T cells (ATCC) and RAW 264.7 cells (ATCC) were maintained in DMEM, 10 % FCS, 100 U/ml penicillin/streptomycin, 2 mM L-glutamine. G418/neomycin (500 µg/ml, *Cambridge BioResources*) was added to transfected cells (TLR8/TFEB-expressing HeLa, TLR8-expressing RAW 264.7).

### Plasmid constructs

TLR8 was cloned into a pcDNA3.1 vector (adding c-myc or V5 tags where indicated) and the TLR8 M1V variant generated by site-directed mutagenesis (QuikChange® II XL; *Stratagene*). TLR8/2 chimera constructs were generated by PCR cloning to fuse the transmembrane (TM) domain of TLR2 with the extracellular TLR8 domain using the Platinum® Taq DNA Polymerase High Fidelity Master Mix (*Invitrogen*) with TLR8/2 primers. The primers used were specific for TLR8 amino acid residues 1-843, namely, TLR8 Forward 5’-CAGAAACATGGAAAACATG TTCCTTCA GTCGTCAATGC-3’; TLR8-2 Reverse 5’-CACATGCCAGACACCAG TGCTGTCATAACC ATGGTGGTGATAAAGAACG-3’. For TLR2, primers for the transmembrane domain amino acid residues 588-610, TLR8-2 Forward 5’CGTTCTTT ATCACCACCATGGTTATGACAGCACTGGTGTCTGGCATGTG-3’and TLR2-TM Reverse 5’-CCTAGGACTTTATCGCAGCTC TCAGATTTACCCAAAATCC-3’ were used. The fragments were then purified, combined and used as templates for a second PCR with the fragment overlapping sequences and the respective forward and reverse primers. The resultant full-length PCR products were subsequently cloned into pcDNA™3.1/V5-His (Invitrogen).

### THP1 CRISPR knockout library

#### Genome-wide CRISPR library

The CRISPR knockout pooled library plasmids was prepared following the protocol previously described (10). Briefly, the Human Toronto Knockout library (TKO V1; *Addgene* #1000000069) was amplified by transformation in Stbl3 competent cells. Colonies were scraped off plates, pooled and purified using Qiagen endotoxin free Maxi kit. Lentiviral particles were generated by transfecting HEK293T cells with the pooled CRISPR library plasmids, and used to transduce Cas-9-expressing THP-1. Cells were maintained in RPMI 1640, 10% FCS, 100 U/ml penicillin/streptomycin, 2 mM L-glutamine, 1 μg/mL Puromycin (*Invivogen*) and 10 μg/mL Blasticidin (*Thermo Fisher Scientific*). Differentiation into macrophages was achieved by treating cells with 20 ng/mL 12-phorbol 13-myristate acetate PMA for 48h prior to experiment.

After 24h of infection with fluorescent *M. tuberculosis*, THP-1 cells were detached using accutase (*BioLegend*) incubation for 20 min at 37°C 5% CO_2_, spun down 300g 5min and fixed in formaldehyde 4% for 1 hour. Cells were then FACS sorted to obtain the top brightest 21% of the population (together with the total population) using a Sony Biotechnology Synergy High Speed Cell Sorter. Genomic DNA was purified using Qiagen FFPE purification kit for fixed cells and subjected to PCR using the following primers: Fwd 5’-AGGGCCTATTTCCCATGATTCCTT-3’, Rev 5’-TCAAAAAAGCACCGACTCG-3’. PCR products were purified using Qiagen PCR clean up and reamplified using the following primers containing P5/7 adaptors as well as appropriate indexes (showed in brackets) necessary for Illumina sequencing: Fwd 5’-AATGATACGGCGACCACCGAGATCTACACTCTCTTGTGGAAAGGACGAGGTACCG-3’, Rev 5’-CAAGCAGAAGACGGCATACGAGAT [TCACTGT]GTGACTGGAGTTCAGACG TGTGCTCTTCCGATCTATTTTAACTTGCTATTTCTAGCTCTAAAAC-3’. PCR reactions were cleaned up and remaining low molecular weight contaminants removed by AMPure XP beads purification using a ratio of 1.6:1. Purification and quantitation were validated using Agilent DNA 1000 chips and confirmed by qPCR. Sequencing was performed on Illumina Hiseq NGS using a Hiseq Rapid SBS kit V2 50 cycles (Illumina).

#### CRISPR screen analysis

Read counts were quantified using the cluster-based approach CB2 by aligning against the Toronto Knock out Library library (11). Guide counts were normalized to counts per 1M sequencing reads for every sample. The values for the unsorted population from all three experiments were combined, choosing the highest count across experiments to represent each guide and compared to the sorted samples from each of the three independent experiments. An aggregate fold change was calculated conservatively as the minimum fold change between the unsorted population and each of the three experiments. To test for overrepresentation of guides in the sorted vs unsorted population, a permutation test was performed. Three rounds of label permuting were conducted to accurately simulate the quantification process generating a randomized unsorted guide count distribution for each of the three experiments. Guides that had less than 0.1 counts per 1M reads in the unsorted population were removed from the analysis. As before, the aggregate log fold change was calculated as the minimum fold change across the three (permuted) experiments. For every guide, the p-value was then defined as the fraction of the number of guides that had a higher minimum log fold change value in the real than in the permuted dataset, and the total number of guides analysed.

#### GO terms analysis

To visualise the genes of the CRISPR screen, hits with a P value <0.05 were transformed into approved symbols using HUGO Gene Nomenclature Committee (HGNC, https://www.genenames.org). The approved symbols were then entered into Panther tools software to assign Gene Ontology (GO) terms to all the hits (http://www.pantherdb.org). REVIGO enrichment analysis (12) was then used to reduce and visualise GO terms. GO terms are therefore summarised and redundancy is removed. Finally, a Python script for circular visualisation of GO terms (CirGO) was used for graphic representation (13).

#### Network analysis

To investigate the functional connectivity of genes identified in the CRISPR screen, we constructed a network of interactions from the Pathway Commons database (14) and performed network propagation using a random walk with restart (RWR) algorithm (15). RWR is designed to retain local connectivity between genes by restarting the signal diffusion process after a limited number of steps, with a fixed probability determined by a restart parameter (r).We used the implementation of RWR provided in the dnet package of the R statistical computing environment (16), with r = 0.2 and Laplacian normalisation of the adjacency matrix. The −log10(P value) for each of the 19,102 genes in the CRISPR screen were used as starting weights for propagation. To account for the fact that highly connected genes (nodes) tend to receive higher steady state scores via RWR, we performed a permutation test in which the starting weights for genes (−log10(Pvalue)) were randomised and RWR was performed a total of 30,000 times. An empirical P value foreach gene was then calculated as the proportion of permuted steady state scores at least as large as that observed from the CRISPR screen data. Interactions between genes with P values < 0.05 were (n = 928) were used to construct a sub-network from the Pathway Commons database. From the resulting sub-network, interactions between genes in the top six largest significantly enriched GO terms identified through the REVIGO enrichment analysis were visualised

### Individual CRISPR knockout cell lines

For individual single guide RNA (sgRNA) cloning, pairs of oligonucleotides were designed and ordered from Sigma with restriction enzyme-compatible overhangs, separately annealed and cloned into the transient CRISPR plasmid pSpCas9 (BB)-2A-Puro (PX459) (*Addgene*, Plasmid #48139) as previously described (17). For cloning into lentiviral vector, LentiCRISPRv2 (*Addgene*, plasmid #52961) was digested with BsmBI (*Fermentas*), and the linearized vector was gel purified before ligation of annealed guide oligo pairs (18). The constructs were amplified in Stbl3 cells (*Invitrogen*) cells and plasmids were purified using endotoxin-free maxi kits (*Qiagen*). Lentiviral particles were produced by co-transfection of LentiCRISPRv2 constructs, psPAX2 (*Addgene*, plasmid #12260), and pCMV-VSV-G (Addgene, plasmid #8454) at a 1:2:1 ratio into HEK 293ET cells using TransIT-293 Transfection Reagent (*Mirus Bio LLC*) reagent according to manufacturer’s instructions. TLR8 and ATG12 CRISPR knockout in THP-1 cells were generated by cloning relevant targeting guide sequences into lentiGuide-Puro vector (*Addgene*, #52963), producing viral particles by transfection into HEK 293T cells as previously described (18) and subsequently transducing THP-1 cells expressing lentiCas9-Blast (*Addgene*, #52962). Cells were maintained in RPMI 1640, 10% FCS, 100 U/ml penicillin/streptomycin, 2 mM L-glutamine, 1 μg/mL Puromycin (*Invivogen*) and 10 μg/mL Blasticidin (*Thermo Fisher Scientific*). Single cell clones were expanded, sequenced to confirm gene knockout, and then pooled. To stimulate differentiation into macrophages, cells were treated with 20 ng/mL 12-phorbol 13-myristate acetate PMA for 48h prior to experiment.

### Transfections

HEK293T cells, HeLa cells, FLAG-tagged TFEB reporter HeLa cells (9) were transfected using Lipofectamine® 3000 Transfection Reagent (*Invitrogen*) and THP1 cells using Lipofectamine LTX (*Invitrogen*) according to manufacturer’s instructions, and assayed 48h post transfection. Primary human macrophages and RAW 264.7 macrophages were nucleofected using the Amaxa Cell Line Nucleofector Kit V (VCA-1003, *Lonza*) and Nucleofector™II Device with programs Y-010 and D-032, respectively. Prior to transfection, complete cell culture medium was removed, and cells were incubated at 37 °C 5% CO_2_ in DMEM containing 10 % FCS. Cells were evaluated at least 48 hours post transfection, either by western blot or immunofluorescence.

### siRNA experiments

For silencing experiments, Accell SMARTpool siRNA for Human TLR8 was obtained from Dharmacon with target sequences against CAAUUAAUAUAGAUCGUUU, CUGGGAUG UUUGGUUAUA, CUAUCAACUUGGGUAUUAA and GUCUUGACUGAAAAUGAUU. PMA-differentiated THP-1 cells were transfected with 1μM of siTLR8 according to manufacturer’s protocol, and assayed 72h post-transfection.

Primary human macrophages were differentiated for three days and transfected with 1μM of either siTLR8 or other PRRs siRNApools using HiPerFect transfection reagent (*Qiagen*) for 5min, and complexes were added drop by drop onto the cells and incubated for 6hours. DMEM was added afterwards and cells were kept at 37 °C for 3 more days. The following sequences were used: (TLR2) GCAAUAACUACGUUUUCUA, UUGUGACCGCAAUGGUAU, UUCUC AUCUCACAAAAUUG, UCUUUAUGUCACUAGUUAU; (TLR3) GCAGCAU AUAUAAUUC AUG, UCACUAUGCUCGAUCUUUC, UUGGAUGUAGGAUUUAAC, CUGU UAGCCAU GAAGUUGC; TLR4: CUAGCUUUCUUAAAUCUUA, CUCUCUACCU UAAUAUUGA, UUCUGGACUAUCAAGUUUA, CCUAUAAGCUAAUAUCAUA: (TLR7): CUAUCGUGCA UCUAUGAAU, CUGUGAUGCUGUGUGGUUU, CUAUGAUGCUU UUAUUGUG, GUAUCA GCGUCUAAUAUCA; (TLR8) CAAUUAAUAUAGAUCGUUU, CUGGGAUGU UUGGUUU AUA, CUAUCAACUUGGGUAUUAA, GUCUUGACU GAAAAU GAUU; (TLR9) GCCACAACU UCAGCUUCGU, CUUGGAUCUGUCACGGAAC, CCUUCGUGGUCUUCGACAA, CCUGC AAAUACUAGAUGUA; (NOD1) CCUUCGUCCUG CAUCACUU, CGUUCAGGUC GAAAGCUUC, GGCUUAUCCA GAAUCAGAU, CCAGCAGUCCUAUGAGUUU; (NOD2) GCCACAUGCAAGAAGUAUA, CGAGCAAUUGCAGAAGUUA, GCUUUAGGA UGUACA GUUA, CUGUUAACCUUU GAUGGCU; (RIG-I, DDX58): GUAUAAUCAUGGAUCGCUU, GGAUUAGCGACAAAUUUAA, CUGACAUACAGAUUUUCUA, UCUUGAUGCGU CAGUGA UA; (Non-targeting Control) UGGUUUACAUGUCGACUAA, UGGUUUACAUGUUUUCUGA, UGGUUUACAUG UUUUCCUA, UGGUUUACAUGUUGUGUGA.

### Mycobacterial infections of macrophages

Infection of primary human macrophages was adapted from Schiebler *et al*. (3). Primary human macrophages WT or knocked down with siTLR8 were infected with *M. tuberculosis H37Rv, M. tuberculosis* Bleupan, *M. bovis* BCG or *M. bovis* BCG-lux at a multiplicity of infection (MOI) of 5:1 for 2 hours, washed in PBS and incubated at 37°C for 24 hours. At indicated time points cells were washed repeatedly, lysed in ddH_2_O, serially diluted and plated onto Middlebrook 7H11 agar plates for CFU enumeration or cell-associated luminescence measurement. Infection of THP-1 macrophages with *M. tuberculosis* Bleupan, *M. bovis* BCG, or clinical isolates of drug sensitive and drug resistant *M. tuberculosis*, was performed at a MOI of 5:1. Infected macrophages were harvested at defined time points, lysed in ddH_2_O, serially diluted, and plated on 7H11 agar medium supplemented as described above.

For infection of THP-1 cells with WT, VirR-, VirR-:WT strains, bacteria cultures were grown as previously described, harvested at mid-log phase, resuspended in PBS and fluorescently labelled with Carboxyfluorescein succinimidyl ester (CFSE) kit (*Biolegend*, 423801) for 30 minutes at 37°C. Bacterial suspensions were then washed twice in PBS with centrifugation steps (3000 g for 10 min) to remove supernatants, and bacterial pellets were finally resuspended in DMEM, 10% FCS prior to macrophage infection. Infection was carried out as described above and cells were incubated either with or without 200 μg/ml RNAse A (*Qiagen*) for 2 hours prior to fixing and immunostaining for MyD88.

### Confocal imaging

Immunofluorescence experiments were undertaken as previously described (3). Cells were seeded on glass coverslips in 24-well tissue culture plates prior to infection with either *M. tuberculosis* H37Rv or *M. tuberculosis ΔleuD ΔpanCD* (Bleupan) double auxotroph expressing either GFP-or mCherry. Following incubation at various time points, cells were washed with PBS, fixed wih 4% paraformaldehyde (PFA) in PBS for at least 30 min and permeabilized for 5 minutes with 0.1% Triton X-100 prior to immunostaining. Primary antibodies (against MyD88, V-ATPase, Ubiquitin, NDP52, LC3, V5 or Myc) were diluted to recommended concentrations in staining medium (DMEM, 10% FCS, 10 mM Glycine, 10 mM HEPES pH 7.4) to which IgG (1:00) was added and the cells were incubated at room temperature for 2 hours. Cells were subsequently washed twice in staining medium. Cells were incubated with secondary antibodies (Alexa Fluor 555 and 647 (*Invitrogen*)) for 30 minutes, protected from light. The cells were subsequently washed, and the coverslips were dipped in water prior to mounting on slides using ProLong™ Gold Antifade Mountant with DAPI (*Thermo Fisher Scientific*). Slides were left to dry overnight, protected from light. Images were acquired either on a Zeiss LSM780 or LSM880 confocal microscope (Plan-Apochromat 63x/1.40 Oil immersion lens) and analysed with Zen 2010 software, Zeiss LSM Image Browser (*Carl Zeiss*), or NIS Elements AR analysis (*Nikon*) software and ImageJ.

For analysis of lysosomal number and acidification in THP-1 macrophages, uninfected THP-1 macrophages were either treated with the TLR8 agonist R848 at 10 μg/ml or left untreated for 24 hours at 37°C, subsequently washed and incubated with 40 nM LysoTracker® Red DND-99 and 1 μM LysoSensor Green DND-189 (*Thermo Fisher Scientific*) for 15 minutes. The cells were then washed twice with PBS after which Live Cell Imaging solution (*Invitrogen*, A14291DJ) was added. Live confocal imaging was carried out on the Zeiss LSM 780 UV. Quantitation of lysosomes was performed using the ImageJ plugin on Fiji app. HEK 293T co-transfected with TLR8 WT - Myc tagged and TLR8 M1V-V5 tagged were fixed in methanol-acetone for immunofluorescent staining with anti – Myc (*Invitrogen*) and anti - V5 antibodies, and counterstained with Alexa 488 and Alexa 647 conjugated secondary antibodies. All cells were visualised using a Leica True Confocal Scanner SP5.

Bone marrow-derived macrophages were generated (as previously described (7)) from femurs of transgenic mice stably expressing mRFP-GFP-LC3 (19) (kind gift from Dr David Rubinsztein, Cambridge, UK), and were either left untreated or treated with 10ug/mL of TLR8 agonist R848 for 24hours at 37°C 5% CO_2_. Cells were then washed with PBS, and incubated in live cell imaging solution (*Invitrogen*) prior to live confocal imaging to visualize lysosomes. Imaging was carried out using on a Zeiss LSM 780UV microscope and quantitation of lysosomes was performed using Image J.

### Cytokine analysis

Primary macrophages were infected with either *M. tuberculosis* ΔleuD ΔpanCD (Bleupan) double auxotroph or *M. bovis* BCG, and either left untreated or treated with R848 10μg/ml. Cell culture supernatant was collected 24 h post infection and analysed using Bio-Plex 17-plex (IL-1β, IL-2, IL-4, IL-5, IL-6, IL-7, IL-8, IL-10, IL-12 (p70), IL-13, IL-17, G-CSF, GM-CSF, IFN-γ, MCP-1 (MCAF), MIP-1β and TNF-α) cytokine assay kit (*Biorad*) according to manufacturer’s instructions.

### Activation assays to determine TLR ligand

For Dual luciferase assay, HEK293T cells were co-transfected with TLR8/2 and the pGL4 luciferase reporter vectors (*Promega*) using Geneporter 2 (*Genlantis*) transfection reagent according to manufacturers’ protocol. HEK293T cells expressing the TLR8/2 and the pGL4 luciferase reporter vectors were treated with various ligands 24h post-transfection. Cells were lysed in passive lysis buffer and lysates were analysed for luciferase activity using the Dual-Luciferase® Assay system (*Promega*). The resultant TLR8/2 chimera generated to promote stable surface expression of TLR8 in HEK293T cells were treated with either whole or lysed mycobacteria, TLR8 ligands CL075 or ssRNA40 (*InvivoGen*) or TLR2 ligand Zymosan (*InvivoGen*) to monitor NF-κB signalling by luminescence.

For quantification of NF-κB activation in THP-1 cells, differentiated TLR8 knockdown (or control) THP-1 BLUE cells were infected with *M. tuberculosis* H37Rv, *M. bovis* BCG, *M. marinum, M. scrofulaceum, M. fortuitum*, or *M. chelonae*, or treated with the TLR8 ligand ssRNA40. Supernatants were collected after 24hours and the levels of NF-κB-induced SEAP were quantified by colorimetric analysis, according to manufacturer’s instructions.

### Phylogenetic analysis

Maximum likelihood phylogenetic tree of all isolates tested were constructed using RAxML (version 8.2.8), generated by mapping detected SNP positions to *M. tuberculosis H37Rv* strain. Representative strains from the main six *M. tuberculosis* lineages described by Comas *et al*., (2013) (20) were included in the analysis for genomic context.

### Quantitation of lysosomal degradative capacity

Uninfected control or R848-tretated THP-1 macrophages were incubated at 37°C for 24hours, washed and incubated with Magic Red™ Cathepsin B Kit (*Bio-Rad*) for 1 hour according to manufacturer’s instructions, then washed twice in PBS. Cells were then resuspended in colorless live imaging solution (*Invitrogen*) and transferred onto 96 well plate for detection of cell-associated fluorescence on the CLARIOstar Plus Multi-mode Microplate Reader (*BMG Labtech*).

### Analysis of phagosomal pH and size

The pH of phagosomes containing MTB was assessed as previously described (21). Briefly, primary human macrophages from individuals that were either homo/hemizygous for the WT and M1V alleles were incubated with PFA-killed *M. tuberculosis H37Rv* double-labelled with acid quenchable (FITC) and pH-resistant (Alexa 633) for a 1h pulse and 23 h chase. The cells were analysed by flow cytometry and intracellular calibration was performed as previously described (21).

For electron microscopy visualization, primary macrophages were infected with *M. bovis* BCG (MOI 10:1). After 24 hours of infection, macrophages were washed and fixed in 0.4% glutaraldehyde for 2 hours at room temperature. Samples were then post-fixed in 1% osium tetroxide followed by dehydration in an ascending graded series of ethanol and embedding in LR white resin. Ultrathin sections (50-70nm) were stained with 2% uranyl acetate and lead citrate and examined in a JM1010 electron microscope (*JEOL*). Phagosome area was measured using ImageJ.

### Western blotting

At the indicated time points, cells were washed twice with PBS, and lysed using RIPA buffer (*Sigma-Aldrich*) containing a proteinase inhibitor cocktail and phosphatase inhibitors (*Roche*). Total protein content was quantified by BCA (*Thermo Scientific*) prior to loading at 20μg and resolving on 17% SDS-PAGE gels, and electro-blotting on to PVDF membranes (*Millipore*) in a wet transfer Cell (*Bio-Rad*). PVDF membranes were blocked by incubation in PBS supplemented with 5% (w/v) fat-free milk powder and 0.005 % (v/v) Tween-20 (*Sigma-Aldrich*) for 1 hour at room temperature. Membranes were washed repeatedly and incubated with primary antibodies following manufacturer recommended concentrations overnight at 4°C. Membranes were then washed and incubated for 1 hour with 1:50 000 dilution of the horseradish peroxidase-(HRP) conjugated secondary antibodies: HRP (*Santa Cruz Biotechnology*). Membranes were revealed using ECL Advance™ Western Blotting Detection kit (*GE Healthcare*) according to the manufacturer’s instructions.

### Mouse infection experiment

Specific-pathogen-free female C57BL/6 mice, from 6 to 8 weeks old, were purchased from the Jackson Laboratories, Bar Harbor, Maine. Mice were maintained in the Biosafety Level III animal laboratory at Colorado State University, and were given sterile water, mouse chow, bedding, and enrichment for the duration of the experiments. The specific pathogen-free nature of the mouse colonies was demonstrated by testing sentinel animals. All experimental protocols were approved by the Animal Care and Usage Committee of Colorado State University. The CSU animal assurance welfare number is A3572-01.

C57BL/6 mice were challenged by low-dose aerosol exposure with *M. tuberculosis* using a Glas-Col aerosol generator calibrated to deliver 50–100 CFU of bacteria into the lungs. Information regarding preparation of bacterial stocks and growth characteristics of the various bacterial strains (*n* = 5) used were as previously described. Strain MDR-TB M10 (resistance profile: Low-level fluoroquinolone resistance, Isoniazid, Rifampicin, Ethambutol, Streptomycin and Pyrazinamide) was originally provided by Dr. Chan, (Seoul, Korea). Strain MDR-TB TN5904 (resistance profile: INH (R, 1.6), EMB (S), RIF (R>8), STR (R, 10), KAN (S) was originally provided by B. N. Kreiswirth, (Public Health Research Institute TB Center, Newark, NJ).

On Day 1 after infection, enumeration of bacteria was performed on two mice. Treatment was started from Day 20 to Day 50 after infection and consisted of the following groups: Control (saline; 0.1 ml intraperitoneal injection once daily) and R848 (2mg/kg by 0.1 ml intraperitoneal injection once daily). On days 20, 35 and 50 following infection, bacterial loads in the lungs and spleen, lung and spleen histology, and flow cytometry were determined in 5 mice from each group. Bacterial counts were determined by plating serial dilutions of homogenates of lungs on nutrient 7H11 agar and counting colony-forming units after incubation at 37°C. Histology and flow cytometry analysis was performed as previously described. All experimental protocols were approved by the Animal Care and Usage Committee of Colorado State University, and experiments were performed in accordance with NIH guidelines. To minimize bias, two groups of independent researchers performed the experiment. One group dosed the animals, whereas the second group determined bacterial burden in the different organs. Lesion scores were determined in a blinded fashion. At days 1, 20, 35 and 50 following infection, bacterial loads in the lungs, lung histology, and mononuclear and lymphocytic cellular expressions were determined as previously described. Bacterial counts were determined by plating serial dilutions of homogenates of lungs on nutrient 7H11 agar and counting colony-forming units after 30 days incubation at 37°C. A total of five animals were infected for each time point.

The accessory lobe of the lung from each mouse was fixed with 10% formalin in phosphate buffered saline (PBS). Tissue sections were stained using haematoxylin and eosin and acid-fast stains as previously reported. For *in vivo* experiments, log 10 protection values of at least 0.60 were considered to indicate activity. Statistical analysis was performed by first converting CFU to logarithmic values and evaluated by a one-way ANOVA followed by a multiple comparison analysis of variance by a one-way Tukey test (SigmaStat software program). Differences were considered significant at the 95% level of confidence.

### Statistical analysis

P-values for assays were determined using two-tailed Student’s t-test or ANOVA (as indicated) using GraphPad. Unless otherwise indicated, experiments were performed on at least three separate occasions with at least triplicate samples for each condition and represented as mean and standard error (SEM).

## Notes

### Competing Interest Statement

The authors have declared no competing interest.

